# Impact of an intra-abdominal cooling device during open kidney transplantation in pigs

**DOI:** 10.1101/568402

**Authors:** Alban Longchamp, Raphael P. H. Meier, Nicola Colucci, Alexandre Balaphas, Lorenzo Orci, Antonio Nastasi, Grégoire Longchamp, Solange Moll, Antoine Klauser, Manuel Pascual, François Lazeyras, Jean-Marc Corpataux, Leo Bühler

## Abstract

**Background:** Kidney transplantation using deceased donors still suffers from high post-operative dysfunction rate. During implantation into the recipient, the kidney rewarms. This second warm ischemia time, which is not monitored, is harmful especially if prolonged. We recently developed an intra-abdominal cooling device that efficiently prevents kidney rewarming during robotic transplantation, and prevent ischemia-reperfusion injuries. Here, we tested the benefits of this cooling device during open kidney transplantation in pigs.

**Materials:** Kidneys were procured from large pigs by open bilateral nephrectomy. Following procurement, kidneys were flushed with 4°C Institut Georges Lopez-1 preservation solution, and placed on ice for 128.5 ± 23.2 min. The cooling device was used to continuously cool down the kidney during the vascular anastomosis time.

**Methods:** Animals underwent double sequential autologous open renal transplantation with (n = 7) and without (n = 6) intra-abdominal cooling. Renal cortex temperature and urine output were monitored. The severity of the ischemia reperfusion lesions was analyzed by histology (modified Goujon score).

**Results:** Mean anastomosis time was similar between groups (43.9 ± 13 min). At reperfusion, the renal cortex temperature was lower in the group with cooling (4.3 ± 1.1°C vs 26.5 ± 5.5°C *p <0.001*). The cooled kidneys tended to be protected from injury, including some histopathological ischemia–reperfusion lesions. With the device, kidneys had a better immediate post-operative urine output (*p=0.05*).

**Conclusions:** Our results indicate that the intra-abdominal cooling device significantly reduces second warm ischemic time during transplantation, is technically safe, and does not prolong anastomotic time.

## Introduction

Kidney transplantation improves survival, is cost effective, and offers the highest quality of life to patients with end-stage renal disease [1-3]. Kidney transplantation outcomes are variable, and depend on non-modifiable factors such as donor and recipient age, pre-existing diseases, or time on dialysis of the recipient, as well as modifiable, such as graft warm ischemic time [4]. The rate of post-operative renal dysfunction has increased over the last years for the transplants performed with deceased donors, in correlation with a higher use of marginal donors [5].

The current consensus is that longer ischemia time primes the tissue for the generation of oxygen reactive species (ROS), and damage upon reperfusion [6]. During ischemia, adenosine triphosphate (ATP) decreases, initiating the dysfunction of mitochondrial ion channels. Dysfunction of mitochondrial ion channels/transporters mechanism, leads to an increase in mitochondrial inner membrane permeability, alteration of anti-oxidative capacity, mitochondrial death and production of excessive ROS. Ultimately, these perturbations during ischemia render cells more susceptible to oxidative stress at reperfusion [7] [8]. While considered unspecific for decades, ROS production during reperfusion was recently demonstrated to occur via reverse electron transport at mitochondrial complex I [9]. During ischemia succinate (citric acid cycle intermediate) accumulates [9], which is rapidly re-oxidized by succinate dehydrogenase during reperfusion to generate large amount of ROS. Thus, reducing mitochondrial energy expenditure in condition of low oxygen and nutrient availability, to maintain ATP homeostasis are of great interest to limit IRI.

Hypothermia decreases metabolic rate, and preserve ATP levels. In kidneys, temperatures below 18 °C prevent from ischemic damage [10, 11]. Prolonged second warm ischemic time (defined as the time during the implantation) is a risk factor for delayed graft function [12]. Similarly, longer anastomosis times are associated with a higher rate of delayed graft function, acute and chronic rejection, and a shorter graft survival [13, 14]. In human, anastomosis time above 45min independently increases the risk of delayed graft function, and impairs allograft function 3 years after transplantation [15]. Importantly, increasingly used kidneys from marginal donors are less resistant to warm ischemia [16]. Any delayed function has important implications, as it prolongs length of stay, increases the need for dialysis, and adversely affects long-term survival [17-19].

Thus, to minimize ischemic time and reduce IRI, efforts have been devoted to maintain a cold environment during the different steps of transplantation. These include the development of complex perfused transport systems, used during the pre-implantation sequence. However, during the intra-operative sequence, temperature is not monitored, and surgeons typically use intermittently splashing of the kidney with cold saline or microparticulated ice. More advanced methods involves immersion in bags, stockinettes or net holding ice with, or without cold preservation chemical [20]. Of importance, none of these methods can achieve a controlled or homogenous cooling [21]. Moreover, none of these methods have been translated to daily clinical practice, this despite encouraging results.

Recently, we developed a cooling device, made of double silicone device in which ethanol and methylene blue continuously circulate. In the setting of robot-assisted kidney transplantation, this device, allowed strict temperature control, which prevented kidney ischemia-reperfusion injuries, [22]. During standard open kidney transplant, vascular anastomoses are easier, and faster, thus second warm ischemic is relatively short. Therefore, in the setting of open transplantation, we questioned if our cooling device was similarly efficient at reducing IRI and improving short-term outcomes.

## Material and methods

### Animals

The study was approved by the University of Geneva animal ethics committee (protocol number: GE/53/14/22826). Five-month-old female pigs (n = 13) with an average weight of 44.7 ± 2.3 kg were obtained from the animal facility of Arare, Switzerland. All pigs were maintained under standard conditions. Water and food were provided *ad libitum*. Animals were allocated into 2 groups: conventionally performed open auto-transplant without (Ctrl, n = 6), or with (Cooling, n=7) cooling system.

### Surgery

Animals were first premedicated using azaperone (2.2 mg/kg IM), midazolam (1.6 mg/kg IM), and atropine (0.02 mg/kg IM), and anesthetized with ketamine (2–6 mg/kg/h), fentanyl (4–6 µg/kg/h), midazolam (0.2–0.4 mg/kg/h), and atracurium (1 mg/kg/h). Animals were then intubated, and ventilated before a nasogastric tube was placed. The arterial line was placed in the internal carotid artery. Monitoring included heart rate, systemic blood pressure, pulse oximetry, and end-tidal CO_2_.

Surgery was performed as described [22]. Briefly, following a midline incision, the peritoneal cavity was opened, and the bowels were reclined. First, the aorta, vena cava and renal vessels were prepared. The pigs received 300 UI/kg heparin intravenous injections. Kidney temperature was measure every 10sec using a thermal probe. Kidneys were explanted and instantly flushed with 4°C Institut Georges Lopez-1 preservation solution. After 1hrs on ice, both kidneys were transplanted sequentially onto the vena cava and aorta using 6-0 running suture. In the cooling system group, the device surrounded kidney from the back table to the kidney reperfusion *in-vivo*. Pigs were maintained under general anesthesia for 2hrs after kidney reperfusion, and urine output was recorded. After 2hrs, pigs were sacrificed using 100 mEq of potassium chloride (KCl) intravenous injections

### Cooling system

The cooling system previously described elsewhere [22], and was made of a double silicone sheath, continuously perfused with 4°C ethanol and methylene blue (Fig. 1C). External and internal thickness was 5 mm, and 0.8 mm respectively.

**Fig. 1.**
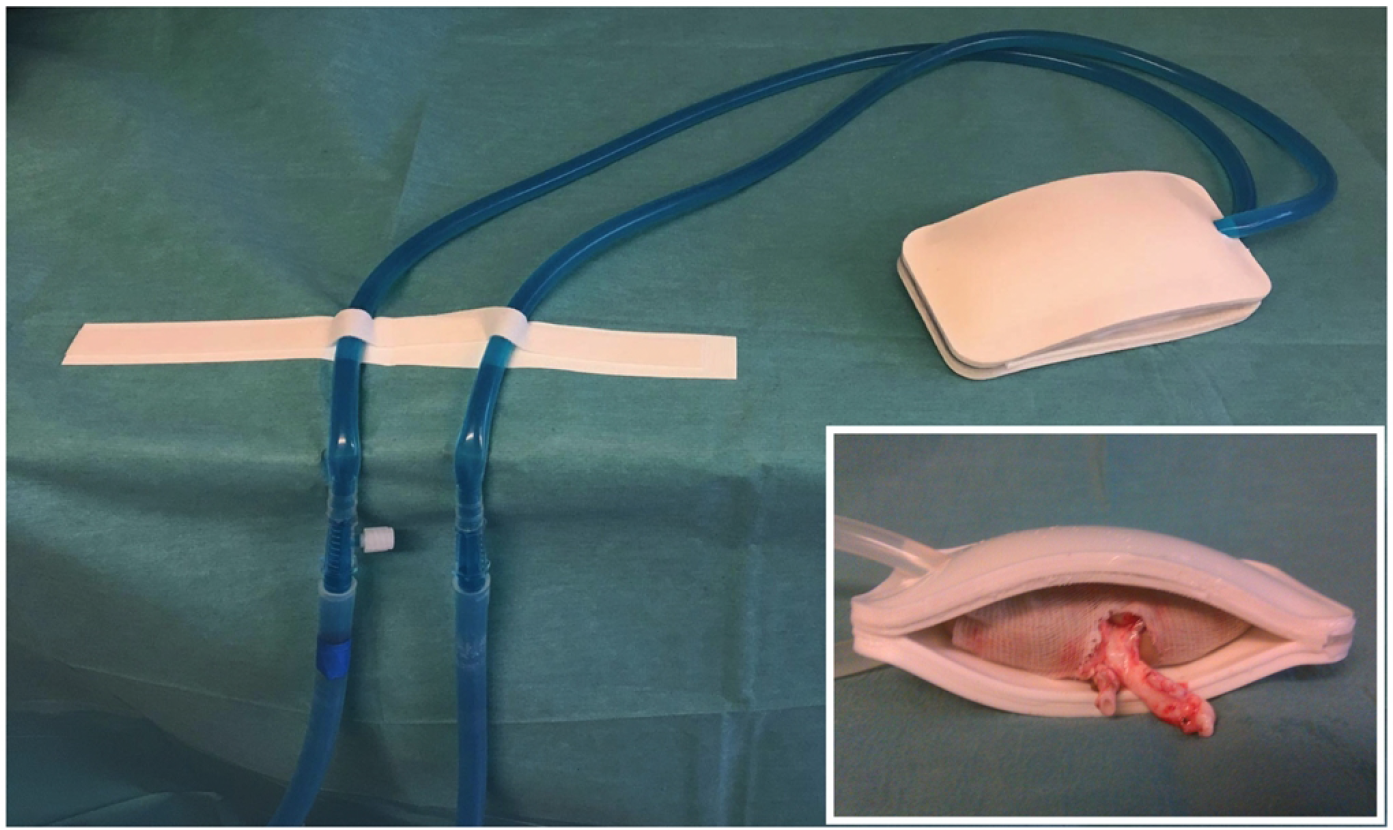
Cooling device. The cooling device with watertight double sheath in silicone surrounding the kidney and continuously perfused by a tubing system with ethanol and methylene blue at 4°C. Enlarged picture: Kidney with the perfused cooling device.

### Histopathological analysis of biopsies

Sections of 3μm thickness were prepared from formalin fixed kidney biopsies, and stained with silver Jones and Periodic Acid-Schiff (PAS). Histopathological analysis score was performed based on those described by Goujon et al.[22, 23] using Osirix software. Four different representative fields were assessed and blinded to group assignment. Lesion severity was graded 0 to 5 according to the following criteria: no abnormality (0), mild lesions affecting respectively 1–10% (1), 10–25% (2), 25–50% (3), 50–75% (4) and >75% (5) of the sample surface. The final score for each biopsy ranges from 0 to 40. A higher score corresponding to the more severe ischemia.

### Statistical analysis

Results were expressed as mean values ± standard deviation (SD). Differences between groups were analyzed using two-sided Student t-test or Mann-Whitney U test. A p value <0.05 was considered statistically significant. Computations were performed using Prism 7 (GraphPad Softwares, San Diego, CA, USA).

## Results

We tested the potential of a cooling device (Fig.1), to reduce second warm ischemia time during vascular anastomosis, and protect from IRI. The cooling device was connected to a tubing system allowing a continuous circulation of ethanol and methylene blue at 4°C. In the open surgery group, the kidney temperature was poorly controlled, reaching 26.5 ± 5.5°C before reperfusion. On the other hand, the cooling device maintained the temperature at reperfusion at 4.3 ± 1.1°C (*p<0.001*, Fig. 2). Similarly, the temperature slope from the start of the anastomosis to the reperfusion was significantly higher in the cooling group (Fig. 2). The temperature drop during flushing, and the temperature on the back table (static, cold ischemia) was similar between the two groups (Fig. 2). Overall operative time was similar, and surgery lasted 366.2 ± 40.5 min (Fig. 3A). Similar surgical technique was used, and the time between kidney artery clamping and flush with cold IGL-1 was 3.5 ± 1.9 min (Fig. 3B). To mimic clinical scenario, after cold flushing the kidneys were kept on ice for 128.5 ± 23.2 min (Fig. 3C). Of importance, the device did not prolong the anastomosis time (43.9 ± 13 min in both group, Fig. 3D) nor increased technical difficulties.

**Fig. 2.**
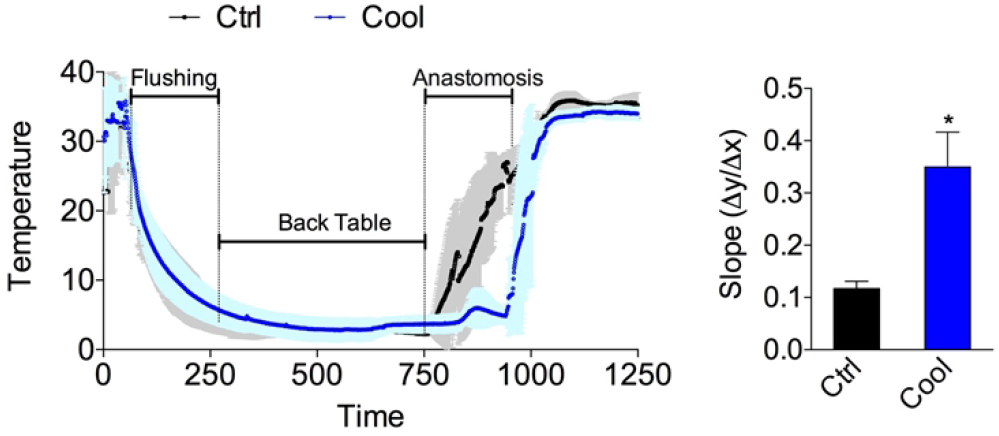
Temperature curve from procurement to reperfusion. Curve (left) and slope during anastomosis (right) in the control or cooling group. n=6-7. Errors bars indicate SEM. \**p*<0.05.

**Fig. 3.**
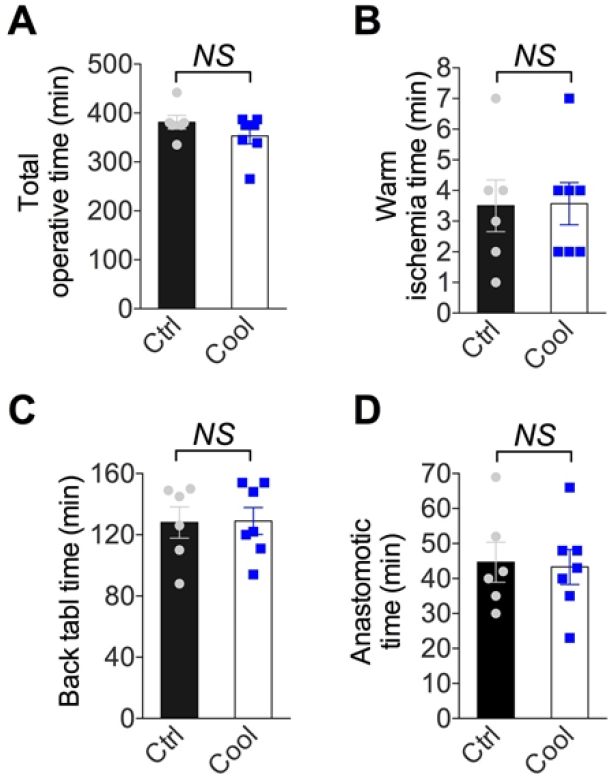
Intra-operative data. (A) Total operative time. (B) warm ischemia time. (C) back table time. (D) Anastomotic time from control and cooling kidney. n=6-7. Errors bars indicate SEM. NS *p*>0.05

Following ischemia and reperfusion, kidneys are at risk of delayed function, characterized by a reduction in urine output. On a cellular level, IRI induces glomerular floculus retraction, brush border loss and tubular dilatation. Ischemia-reperfusion injuries were slightly reduced in the group with cooling, compared to without (Ctrl, Fig. 4), but this failed to reach significance. In cooled group, the percentage of glomeruli with Bowman’s space showing a retraction of the flocculus was lower. However, the amount of cellular debris, brush border loss, and dilated tubule were similar. When these changes were analyzed independently, no significant difference was observed (Fig. 4A and B). The Goujon score [22], combines all the previous histopathological observations into a score, and suggested that cooling protected from IRI (Fig. 4). Urine output within the first days after transplantation defines graft function, and thus allows functional assessment of kidney function following IRI. Within the first hour after reperfusion, urine output was higher in cooled kidneys compared to the control (Fig. 5). This is consistent with a protective effect of continuous, intra-abdominal cooling during transplantation.

**Fig. 4.**
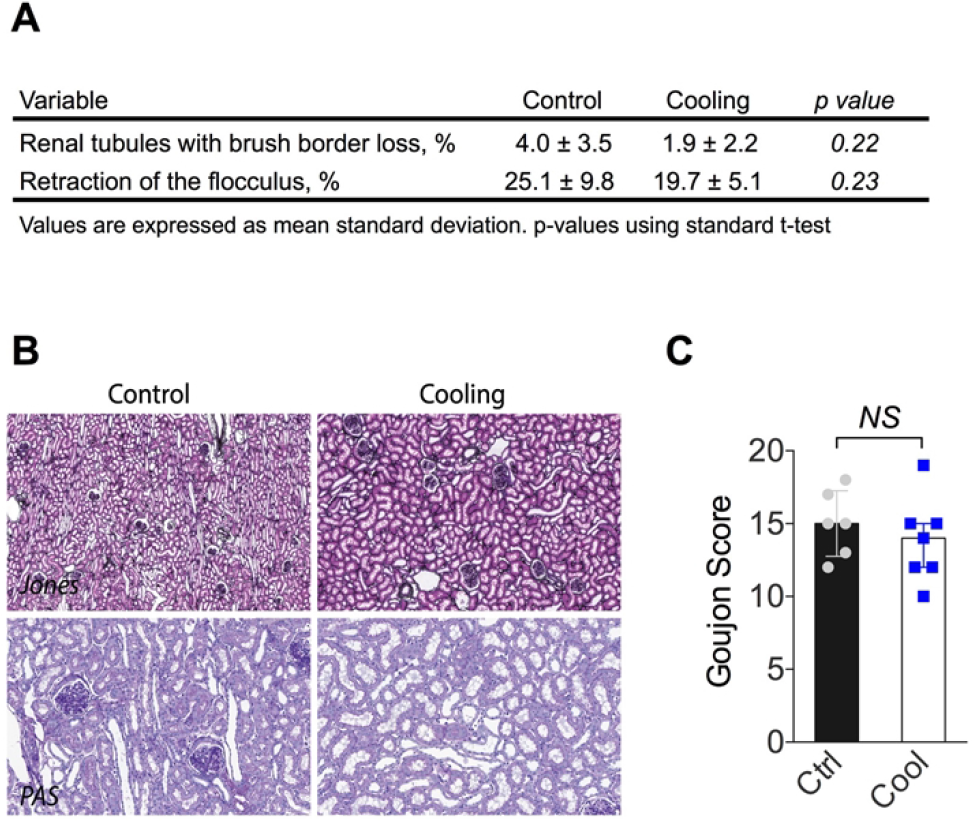
Histological analyses following kidney reperfusion. (A) Percentage of lumina of tubules with cellular debris, the percentage of brush border loss, tubular dilatation, and percentage of floculus retraction in Bowman’s space. (B) Representative kidney histological sections stained with Periodic Acid-Schiff (PAS) (proximal tubular lesions) and Jones (glomerular injury). (C) Goujon score (lesion severity) was graded 0 to 10 for each type of lesion. n=6-7. Error bars indicate SEM. NS *p*>0.05

**Fig. 5.**
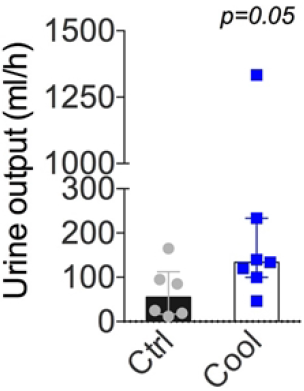
Urine output following transplant. Mean urine output during the first hour following revascularization. Errors bars indicate interquartile range.

## Discussion

Solid organ transplantation would greatly benefit from measures that attenuate the ischemia and reperfusion injury. Here, we present experimental evidence confirming that current clinical implantation methods result in significant rewarming of kidney grafts. The use of our cooling device significantly reduced this rewarming and allowed efficient continuous cooling of the kidney. While short-term histological measures of injury were not statistically different between the groups, cooling of the kidney improved immediate function. Importantly, the cooling device provided continuous, controlled temperature control. The device was safe and did not prolong the surgical time.

Histology of kidneys collected 2hrs after reperfusion, showed consistent ischemia-reperfusion lesions. The most significant lesion was retraction of the floculus, which represented 25.1% and 19.7% of the control and cooling kidneys respectively. In pigs, we demonstrated that during robotic transplant with anastomotic time of 70 min, continuous cooling of the kidney led to a 25% reduction in histopathological damage compared to open standard kidney transplant [22]. In human, warm ischemia time above 60 minutes are associated with a 25% higher risk of death or graft failure [6]. In our study, anastomosis time was similar between groups (43.9 ± 13 min). We believe that the cooling device would provide more benefits in the context of longer anastomosis [22]. Thus, its benefits might depend on factor that complicate the surgery, such as pre-existing vascular diseases, vessel length, or recipient body mass. Conversely, the absence of time constrain might facilitate the teaching of young trainees. Here, total operative, cold and warm ischemia time were similar between the two groups, further suggesting that these differences were due to the prolonged secondary ischemia time during anastomosis. While histological lesions were not statistically different, urinary output and thus immediate function was improved in the cooling group. Altogether, this supports the utility of an efficient and continuous cooling device, especially in human, when a technically demanding anastomosis can be foreseen.

Early graft dysfunction is initiated by IRI [24, 25]. Thus, to ameliorate kidney transplantation, several clinical trials (e.g. peri-operative estrogen administration NCT03663543) are aiming to reduce IRI. It is unlikely that a single intervention suffices. Complementary strategies, in addition to those that reduce exposure to warm ischemia time might have more meaningful benefits. A group that would considerably benefits from shorter warm ischemia time are expanded criteria organs (ECD), defined as organs from donors older than 60 years or 50 years with associated vascular comorbidities [26]. ECD represent half of kidney donors, and are utilized more and more frequently in our aging population [27]. Importantly, ECD are more prone to IRI, and both graft, and patient survival is reduced compared to healthy young donors [28]. It is possible that the slight benefits observed here with a controlled, continuous organ cooling, would have significant impact on patient survival, with the most benefits in the most marginal organs.

Our study has several limitations. We chose an auto-transplant experimental model to avoid alloimmunity, potential inadequate immunosuppressive regimens, toxicity, and acute rejection [29] [30]. Furthermore, the relatively short cold ischemia time corresponded mainly to a living donor situation, where IRI are limited. The current experiments were performed on healthy, young adult pigs, more resistant to stress than human (e.g. IRI). These could explain the absence of significant histological differences. In addition, we only examined short-term outcomes, as pigs could not be kept alive for survival surgeries at our establishment.

In conclusion, our results indicate that the intra-abdominal cooling device significantly reduces warm ischemic time during transplantation, is technically safe, and does not prolong anastomotic time. A randomized, prospective clinical trial, using the cooling device is warranted to evaluate the benefits in human.

## Acknowledgements

This work was supported by the Swiss National Science Foundation to JMC and FL (SNSF 320030_182658), The Mendez National Institute of Transplantation and the Leenards Foundation to AL. We also thank Jean-Pierre Gilberto, and Raphael Ruttiman for their excellent technical assistance.

## Author contributions

AL, RPHM, NC, AB, LO, AN, GL performed the experiments. AL RPHM, NC and LB analyzed the data. AL, RPHM, FL, AK, MP, JMC and LB wrote the manuscript. AL, JMC, FL and LB obtained funding.

